# cDNA-guided functional selection uncovers selective defense systems against RNA phages

**DOI:** 10.64898/2026.04.06.716636

**Authors:** Hee-Won Bae, Hyeong-Joon Ki, Shin-Yae Choi, Han-Gyu Cho, Chae-Hyeon Woo, Min-Ju Kim, Hyung-Jun Chun, You-Hee Cho

**Affiliations:** Program of Biopharmaceutical Science and Department of Pharmacy, College of Pharmacy and Institute of Pharmaceutical Sciences, CHA University, Gyeonggi-do 13488, Korea

**Keywords:** RNA phage, cDNA, defense systems, defense islands, *Pseudomonas aeruginosa*

## Abstract

Bacteria encode diverse antiphage defense systems, yet mechanisms that target RNA phages remain comparatively underexplored. Here, we used a cDNA-based functional selection strategy to systematically identify genes that confer resistance to RNA phage infection independently of receptor variation in *Pseudomonas aeruginosa*. This approach uncovered previously uncharacterized antiphage defense systems, most of which are located within genomic islands, consistent with their being bona fide components of bacterial immune systems. Several systems conferred selective resistance to RNA phages and their carriage was associated with pilin variability, suggesting layered anti-phage immunity. Among these systems, Zws is the most prevalent RNA phage defense system and functions as a multidomain effector. Structural modeling and in vitro cleavage assays showed that ZwsA is an RNA endonuclease that selectively cleaves RNA phage genomes through a predicted NERD domain. Together, these findings expand the current framework of bacterial antiphage immunity and highlight the power of functional genomics to uncover cryptic components of the bacterial antiviral arsenal.

**IMPORTANCE:** Bacteria harbor a broad repertoire of antiphage defense systems, but our understanding of mechanisms that target RNA phages remains limited, being heavily biased toward defenses against DNA phages. By applying a cDNA-based functional selection strategy, this study overcomes a major obstacle in defense-gene discovery and uncovers previously uncharacterized genes that represent bona fide components of the bacterial immune arsenal against RNA phages. The identification and characterization of Zowangsin (ZwsA), a NERD-domain RNA endonuclease that selectively cleaves specific signatures within RNA phage genomes, establish targeted RNA degradation as a central principle of bacterial defense against RNA phages. More broadly, this work expands the conceptual framework of bacterial antiviral immunity and illustrates the utility of functional selection for uncovering cryptic immune systems with implications for phage biology, RNA biology, and biotechnology.

**HIGHLIGTHS:**

- cDNA-based functional screen identifies six defense systems that restrict RNA phages
- These defense systems are enriched within genomic islands of *Pseudomonas aeruginosa*
- Zws, Szs, and Mws systems confer selective defense against RNA phages
- ZwsA is a signature-selective RNA endonuclease that targets phage genomic RNA

## INTRODUCTION

Bacteria have evolved diverse antiphage defense systems to survive in phage-rich environments. Identifying these systems and elucidating their mechanisms are central to understanding the various aspects of phage biology and biotechnology, which enables discovery of new biological phenomena and/or new biotechnological toolkits. For example, abortive infection systems that sacrifice the infected bacterial cells to prevent the spread of progeny phages can be regarded as a social behavior to ensure the survival of the bacterial population [1–3]. Restriction-modification and CRISPR-Cas systems are the antiphage defense systems that have become fundamental tools in molecular biology and genome engineering [4, 5]. Accordingly, increasing attention has been directed toward the discovery of new anti-phage defense systems. Recent studies have shown that antiphage defense systems are often clustered in discrete regions of prokaryotic genomes. [6, 7]. These loci, termed “defense islands”, are rich sources of previously unrecognized defense genes [8, 9] and have enabled “guilt-by-association” strategies for the in silico identification of new antiphage defense systems across microbial pangenomes [10–12]. Many such systems have subsequently been functionally validated in heterologous model bacteria, including *Bacillus subtilis* and *Escherichia coli* [7]. Notably, some genes discovered in defense islands have also provided insights into the evolutionary links between prokaryotic and eukaryotic cellular immunity. For example, prokaryotic viperins, identified through their homology to human viperin, exhibit analogous biochemical and physiological activities [13]. In humans, viperin is induced during infection by DNA and RNA viruses and depletes the cytidine triphosphate (CTP) pool by converting it to 3’-deoxy-3’,4’-didehydroCTP, which results in premature termination of RNA synthesis by viral enzymes [14, 15]. Despite these advances, most antiphage defense studies have focused almost exclusively on DNA phages, whereas defenses against RNA phages remain much less explored, in part because RNA phages are less well characterized and typically require specialized pili for infection [16].

Here, we sought to identify specialized defense systems active against RNA phages, motivated in part by the possibility that some antiviral principles may be conserved across kingdoms. We previously established a cDNA-based reverse genetics platform for the small RNA phage PP7 and used it to investigate key steps in its life cycle during infection of *Pseudomonas aeruginosa* (PA) [17]. As an opportunistic human pathogen and a major model for phage-host interactions, PA possesses a highly dynamic pangenome enriched in defense-associated genomic islands, including two core defense hotspots [18, 19]. In addition, PA displays substantial variation in surface structures such as type IV pili, which mediate phage adsorption [20, 21]. Using this cDNA-based PP7 production platform, we identified six genetically distinct defense systems. These systems are positioned in proximity to known defense modules and/or within core defense hotspots. Functional analyses further showed that several of these loci confer selective resistance to RNA phages rather than broad, nonspecific antiviral activity. Among them, ZwsA displayed signature-selective RNA endonuclease activity that specifically cleaves RNA phage genomes.

## RESULTS

### Six defense systems identified against RNA phages

We previously proposed that cDNA-based RNA phage assembly could be used to identify “intracellular” defense systems against RNA phages by monitoring reduced phage production [22]. Because this strategy bypasses adsorption and genome entry, it excludes defense mechanisms based on infection exclusion, including those mediated by natural pilin variation in PA [16, 23]. As outlined in Figure 1A, we first screened 47 in-house PA strains for reduced production of the RNA phage PP7 from chromosomally integrated cDNA, using the surrogate strain PAK as a reference.

**Figure 1.**
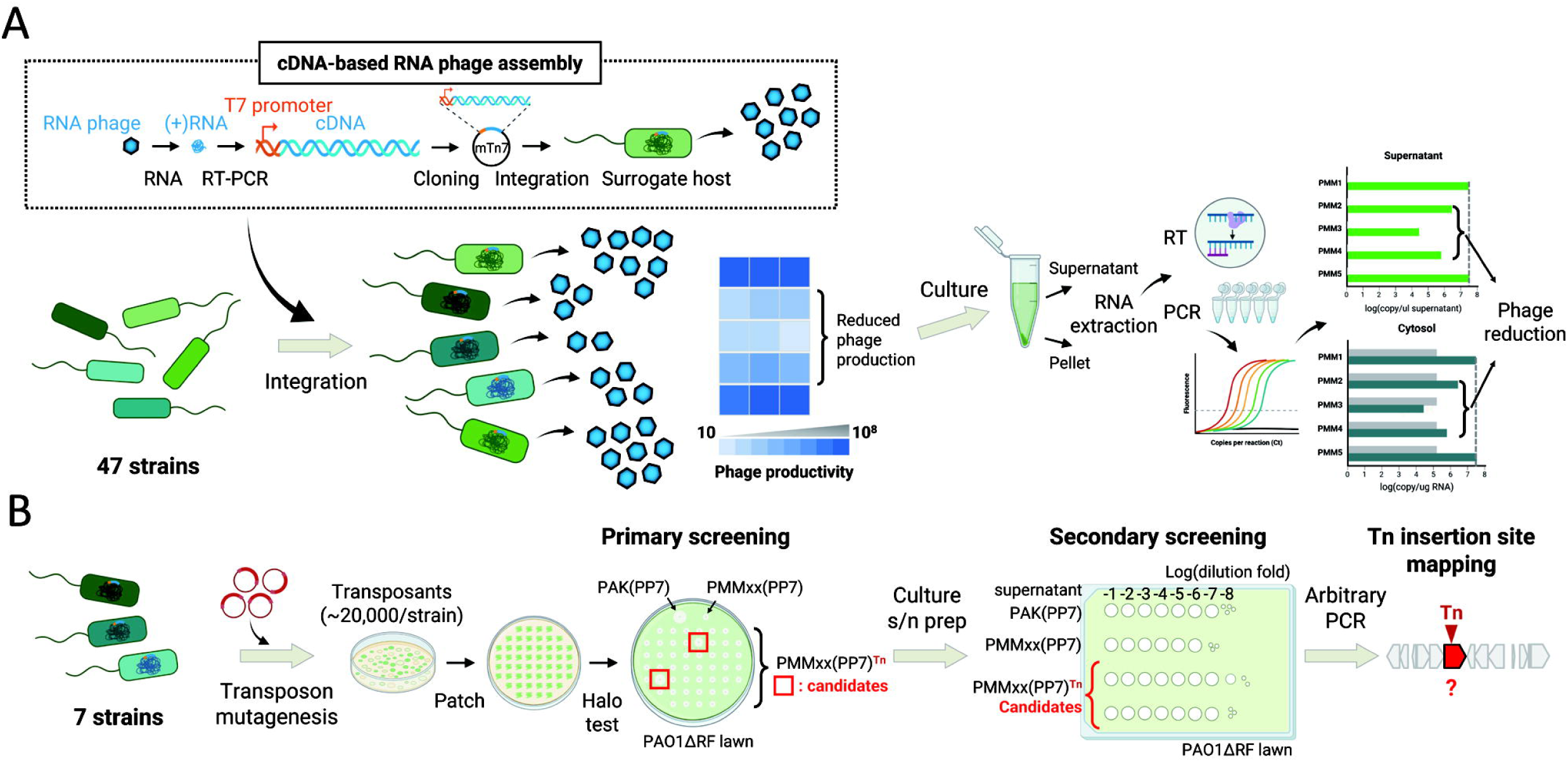
Screening pipeline for identification of bacterial genes that reduce PP7 production. **A**. Schematic workflow for establishing a cDNA-based PP7 production system in the PA strains. The PP7 cDNA has been chromosomally integrated into a panel of the in-house PA (PMM) strains. The PMM strains with the PP7 cDNA are screened for reduced phage production relative to the surrogate host strain, PAK as the control. Strains with decreased PP7 productivity are verified by quantitative analyses for phage progeny production and phage genomic RNA synthesis. **B**. Transposon mutagenesis followed by gene identification. Approximately 20,000 transposants per strain from the selected 7 PA strains carrying PP7 cDNA designated as PMMxx(PP7) (see Figure 2) were tested for enhanced halo formation on a PP7-susceptible strain, PAO1ΔRF, the PAO1 mutant devoid of both R and F type pyocins. Candidate transposants were verified by phage progeny production and then subjected to arbitrary PCR to map the transposon insertion sites responsible for the phenotype.

The first 11 strains exhibiting reduced PP7 productivity were subjected to secondary validation by quantifying genomic RNA in both the supernatant and the cytosol samples. Four of these strains were excluded because of technical issues such as failure in marker excision (PMM13 and PMM18) and irreproducible phage productivity after marker excision (PMM36 and PMM47). The remaining seven strains, which displayed varying degrees of productivity reduction, were subjected to transposon mutagenesis followed by screening for mutants in which phage production was restored to levels comparable to those of PAK (Figure 1B). As shown in Figure S1A and B, PMM14 and PMM51 exhibited the strongest reduction in phage productivity, with decreases of ∼4-5 orders of magnitude, whereas PMM38 and PMM44 showed reductions of ∼3 orders of magnitude. PMM16, PMM19, and PMM55 displayed decreases of ∼2 orders of magnitude. Reduced phage productivity correlated with lower levels of genomic RNA and progeny phage particles, as determined by RT-qPCR (Figures S1C and D). These data suggest that diminished phage production in these strains results from reduced steady-state levels of phage genomic RNA in the bacterial cytoplasm, most likely owing to intracellular bacterial defense activities.

A total of approximately 140,869 transposon insertion mutants were screened across the seven strains, corresponding to ∼20,000 mutants per strain. From this screen, 12 nonredundant mutants were isolated (Figure 2), and insertion-site analysis identified six defense loci (Figure 3). These systems were named after the protective deities in East Asian tradition: *seongzusin* (*szs*; identified in PMM14 and PMM51), *zowangsin* (*zws*; identified in PMM19, PMM38, PMM44 and PMM55), *moonwangsin* (*mws*; identified in PMM44), *teozusin* (*tzs*; identified in PMM38 and PMM55), *obangsin* (*obsC* and *obsG*; identified in PMM16), and *cheolryungsin* (*crs*; identified in PMM44). It is noted that the *szs* genes were identified from strains showing the largest reduction in phage productivity, suggesting that the Szs system may mediate the strongest defense activity among the identified systems. By contrast, the Zws system was independently identified four times across strains showing more moderate reductions, indicating either strain-dependent variation in Zws-mediated defense or a limitation of the screening conditions used in this study. PMM44 is notable that three defense systems (Zws, Mws and Crs) were functionally identified in a single strain, whereas two unlinked defense systems (Zws and Tzs) were functionally identified in both PMM38 and PMM55. In contrast, only one system was identified from PMM14 (Szs), PMM16 (Obs), PMM19 (Zws), and PMM51 (Szs), despite the genomic presence of the Obs and Tzs systems in PMM14 and the Szs and Tzs systems in PMM16 (Figure 3; see below).

**Figure 2.**
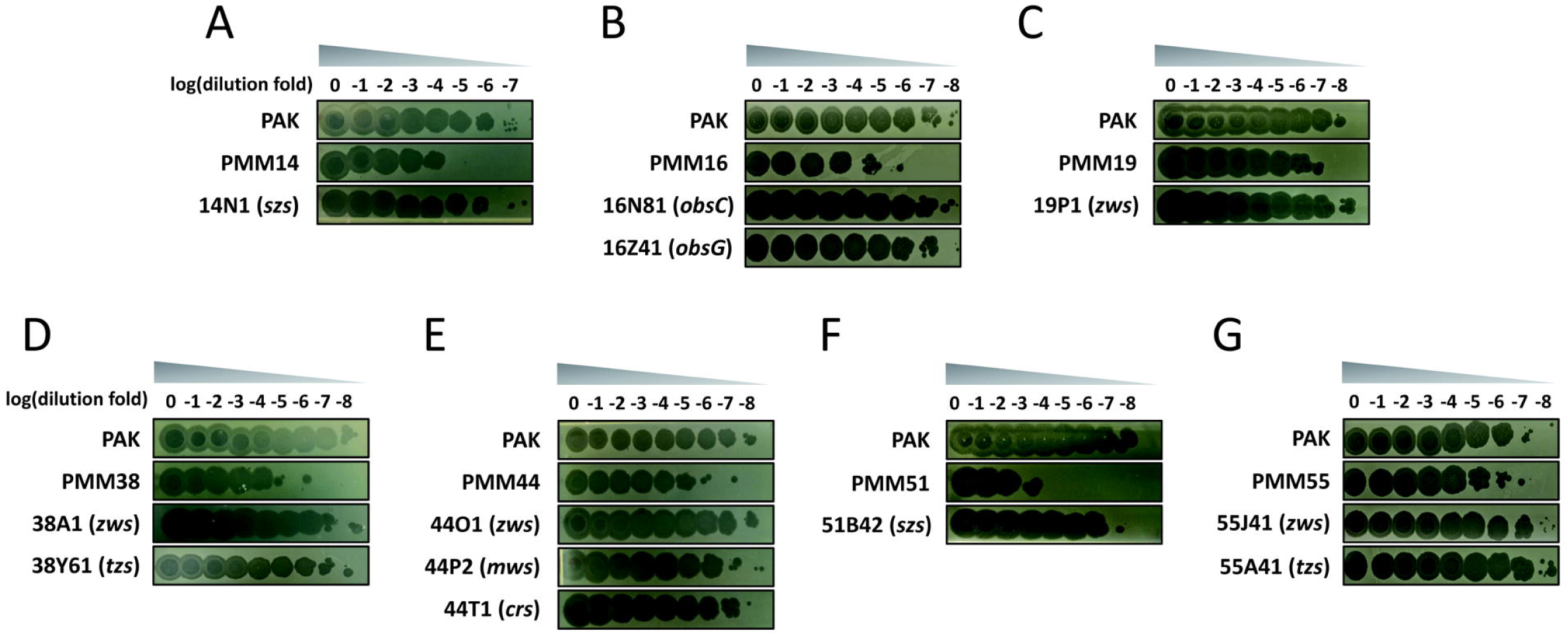
RNA phage productivity of the selected strains and their mutants. PP7 productivity from the 7 selected PMM strains with reduced phage productivity and their 12 transposon mutants isolated as described in Figure 1. Serial dilutions from the 12-h culture supernatants were subjected to spotting assays on PAO1ΔRF, in comparison with those from PAK. The identified gene(s) from each mutant are designated as follows: *szs* for <u>S</u>eong<u>z</u>u<u>s</u>in; *zws* for <u>Z</u>o<u>w</u>ang<u>s</u>in; *mws* for <u>M</u>oon<u>w</u>ang<u>s</u>in; *obsC* and *obsG* for <u>Ob</u>ang<u>s</u>in; *tzs* for <u>T</u>eo<u>z</u>u<u>s</u>in; *crs* for <u>C</u>heo<u>lr</u>yung<u>s</u>in. **A**. PMM14 and 14N1 (*szs*). **B**. PMM16, 16N81 (*obsC*) and 16Z41 (*obsG*) **C**. PMM19 and 19P1 (*zws*). **D**. PMM38, 38A1 (*zws*), and 38Y61 (*tzs*). **E**. PMM44, 44O1 (*zws*), 44P2 (*mws*), and 44T1 (*crs*). **F**. PMM51 and 51B42 (*szs*). **G**. PMM55, 55J41 (*zws*), and 55A41 (*tzs*). Numbers indicate the logarithmic values of the dilution folds.

**Figure 3.**
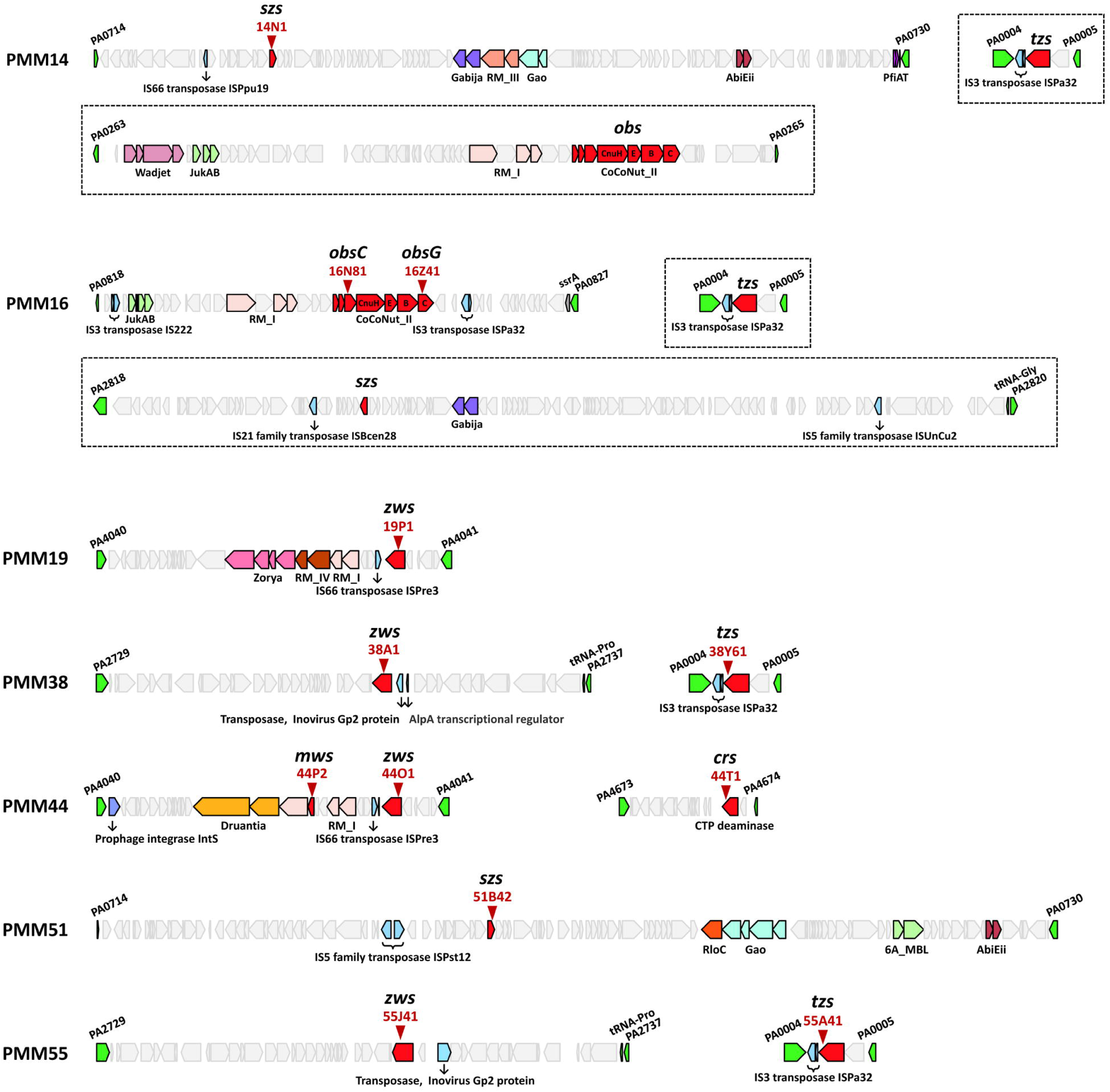
Genetic organization of the identified genes. The 7 selected PMM strains with reduced phage productivity are shown with their gene clusters where the 12 defense systems have been identified from transposon screen. Red arrow heads indicate the insertion sites of the isolated mutants, with the defense genes (red) labeled by the names (see texts). Genes (green) at both ends define the boundaries of the genomic islands. Defense genes identified by DefenseFinder v2.0.0 and mobile genetic elements are colored with the names and non-defense genes (light gray) are shown. The dotted boxes highlight the genes identified from the genome sequences of PMM14 and PMM16.

### Genomic distribution and organization of the identified defense systems

Previous work showed that defense systems in PA are modularly organized at defined genomic loci, particularly within the core defense hotspots cDHS1 and cDHS2 [18]. Known defense systems are frequently clustered within genomic islands, which can be horizontally transferred and thereby promote the acquisition of phage resistance. Such modular clustering is a hallmark of professional or acquired defense systems and is characteristic of accessory genome content. As shown in Figure 3, most of the six systems identified here are located adjacent to known defense genes within defense islands in PMM14, PMM16, PMM19, PMM44, and PMM51, based on the genomic synteny of each strain with either PAO1 or PA14 (Fig. S2A). Although the *zws* loci in PMM38 and PMM55 and the *crs* locus in PMM44 are not directly clustered with previously annotated defense genes, they are also embedded within genomic islands: *zws* lies within a gene cluster at cDHS1 in both PMM38 and PMM55, whereas *crs* is located at the region 42 of genetic plasticity (RGP42) [24]. Together with the co-occurrence of mobile genetic elements, these observations support the conclusion that all the 6 defense systems identified in this study were acquired by horizontal transfer. We also note that *tzs* and *obs* were identified by sequence analysis in PMM14 and that *tzs* and *szs* were likewise identified in PMM16, even though no corresponding transposon mutant was recovered from those two libraries.

Comparative genomic analyses of the 7 strains and PAO1 enabled us to identify the 8 genomic locations for the 14 defense systems from the 7 strains (Figure S2B), which include the already identified core defense hotspots (cDHS1 and cDHS2) and the RGP42 [24]. The 4 genomic positions are the spots for the only one defense system, which include cDHS2 for the PMM16 Obs system. It should be noted that the other 4 genomic positions are for the two defense systems, which include cDHS1. For example, the cDHS1 is for the defense islands harboring the Zws system in PMM38 and PMM55 (Figure 3). The genomic position between PA4040 and PA4041 genes are utilized by the other 2 Zws systems in PMM19 and PMM44, whereas the location between PA0714 and PA0730 genes are occupied by the large defense islands including the Szs system in PMM14 and PMM51. This position has been occupied by filamentous prophage, Pf4 in PAO1, formerly identified as *att1* [25]. It is remarkable that all the Tzs systems are flanked by PA0004 and PA0005 with the same genetic organization with IS3 transposase (Figures 3 and S2B), suggesting that the Tzs system comprises a defense islet between PA0004 and PA0005 from the 4 out of the 7 reduced-productivity strains in this study. This pattern of genomic localization and distribution for the identified defense systems is consistent with horizontal acquisition and modular assembly of the antiphage defense genes within the PA pangenome.

### ZwsA, SzsA, and MwsA confer selective defense against RNA phages

Based on the host ranges of the type IV pilus (TFP)-dependent RNA phages PP7 and LeviOr01, we selected PAO1 and PMM3 from our strain collection for infection by PP7 and LeviOr01, respectively [16, 23]. To test whether the 6 candidate systems indeed confer antiphage activity, each gene or locus was cloned into the multicopy plasmid pUCP18 and introduced into the corresponding phage-susceptible strain. As shown in Fig. 4A and B, multicopy expression of ZwsA, SzsA, and MwsA strongly inhibited plaque formation by PP7 and LeviOr01, but had no detectable effect on the TFP-dependent DNA phages MP29 and MPK7 [26]. ZwsA contains a NERD domain, whereas MwsA contains a PIN-domain RNase module. Both proteins were therefore consistent with candidate antiviral effectors based on domain architecture and genomic association with known defense loci [27, 28]. By contrast, SzsA contains a LOTUS domain, a small, conserved RNA-associated domain originally defined in Limkain, Oskar, and Tudor domain-containing proteins 5 and 7 (TDRD5 and TDRD7) [29]. Because LOTUS-domain proteins are found in both prokaryotes and eukaryotes and often function in RNA-associated complexes for RNA-protein interaction and RNA regulation, SzsA may act as a specialized RNA-binding factor that recruits one or more effectors to target phage RNA.

**Figure 4.**
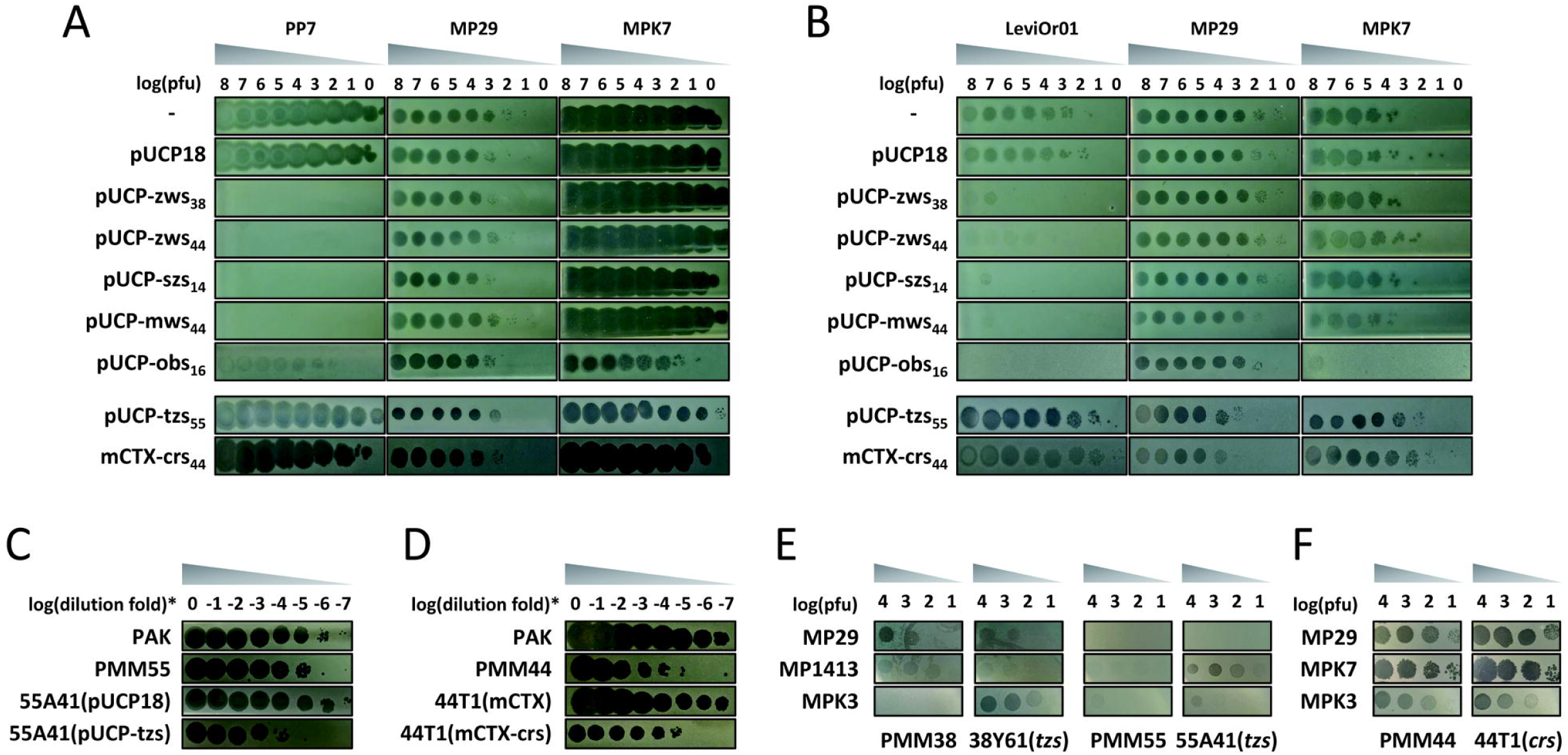
Validation of the defense genes. **A** and **B**. Defense spectra of the defense genes in PAO1 (**A**) and PMM3 (**B**) against phage infections. Ten-fold serial dilutions of phage lysates of two TFP-requiring RNA phages (PP7 and LeviOr01) and two TFP-requiring DNA phages (MP29 and MPK7) are spotted on a lawn of PA cells (-) or those with one of the pUCP18-derived plasmids as designated or with chromosomally integrated gene (mCTX-crs). The numbers in subscript indicate the source PMM strains. pUCP-obs_16_ contains the 7-gene *obs* operon in Figure 3, unlike the others that contain only one coding region. The numbers on top indicate the log(pfu) values. **C** and **D**. Complementation by the defense genes in PMM55 (**C**) and PMM44 (**D**) mutants for the *tzs* gene (55A1) or the *crs* gene (44T1) PP7 productivity was assessed for the mutants with one of the constructs for the *tzs* (**C**) and the *crs* (**D**) genes as in **A**. Serial dilutions from the 12-h culture supernatants were subjected to spotting assays on PAO1ΔRF, in comparison with those from PAK. Numbers indicate the logarithmic values of the dilution folds. **E** and **F**. Defense spectra of the isolated mutants for *tzs* (38Y61 and 55A41) or *crs* (44T1) against phage infections. Ten-fold serial dilutions of phage lysates of DNA phages (MP29, MP1413, MPK3, and MPK7) are spotted on a lawn of PMM strains and their mutants as designated. The numbers in subscript indicate the source PMM strains. Numbers indicate the log(pfu) values.

During this work, we noted the report by Bell et al. [30] which defined a new class of antiviral defense systems termed coiled-coil nuclease tandems (CoCoNuTs). Based on domain organization, CoCoNuTs are predicted to target both RNA and DNA. The *obs* locus from PMM16 appears to comprise seven genes (*obsA* to *obsG/cnuC*) and shows strong homology to type II CoCoNuTs, prompting us to en bloc clone the entire seven-gene cluster into pUCP18. Ectopic expression of this locus protected PAO1 and PMM3 from infection by the RNA phages PP7 and LeviOr01, as well as by the DNA phage MPK7, but not by MP29 (Fig. 4A and B). Unlike RNA phages that inject their genomes through the PilQ ring by detaching the pilin polymers [31], the mechanisms of TFP adsorption and/or genome entry may differ in TFP-dependent DNA phages [22], which accounts for the selective activity of Obs against MPK7 but not MP29 that may reflect differences in the stage at which protection is exerted. Regardless, these data indicate that Obs can protect against RNA phages and at least a subset of DNA phages, consistent with recent predictions for CoCoNuT-like systems. The molecular basis by which the seven-gene Obs system acts against both RNA and DNA phages remains to be determined.

In contrast to the 4 systems described above, ectopic expression of Tzs and Crs did not protect PAO1 or PMM3 from RNA phage infection. However, both loci restored the reduced-productivity phenotype of the corresponding transposon mutants when reintroduced into the PP7-reduced mutant backgrounds (55A41 for Tzs and 44T1 for Crs) (Fig. 4C and D), suggesting that their defensive activities may be strain dependent. Tzs is predicted to encode a DEAD-box protein, a family broadly implicated in RNA- and DNA-associated processes through helicase activity [32]. Given that DEAD-box proteins are conserved across prokaryotes and eukaryotes, it is plausible that Tzs-like proteins participate in antiviral defense, as has been proposed for other antiphage systems such as Hachiman and Druantia [7, 33]. Consistent with a broader defensive role, the isolated *tzs* mutants (38Y61 and 55A41) were more susceptible to the DNA phages MP1413 and MPK3 (Figure 4E), indicating that Tzs may also contribute to defense against DNA phages.

The *crs* locus encodes a protein with homology to nucleotide deaminases. Such proteins are often associated with nucleotide depletion and could therefore restrict phage replication by reducing nucleotide availability. This type of system may also function through abortive infection, with an associated partner mitigating host toxicity [1, 34], although this remains to be tested for Crs. Notably, the isolated *crs* mutant (44T1) did not differ from its isogenic parent in susceptibility to the DNA phages tested (MP29, MPK7, and MPK3) (Figure 4F), suggesting that Crs may act preferentially against RNA phages. Although the precise mechanisms by which Tzs and Crs reduce PP7 productivity remain unresolved, our data support the conclusion that both systems contribute to intracellular immunity against PP7 in their native strain backgrounds.

### Association of the defense systems with the pilin variability

We previously showed that natural variation in pilin sequences can determine susceptibility to specific RNA phages in PA strains, raising the possibility that intracellular defense systems may be shaped by, or compensate for, the lack of pilin-mediated infection exclusion. To examine this relationship, we generated a PilA phylogeny from 653 PA genomes retrieved from the database. As shown in Figure 5, 60.5% of the strains carry G1a (16.8%), G1b (18.2%), and G2c (25.4%) pilins, which associated with susceptibility to LeviOr01 and PP7 [16, 23]. Genome-level analysis further showed that the Zws system is enriched among strains carrying susceptible pilins, particularly G1b pilins (18.2% vs. 42.1%). This is in quite contrast to the defense systems against DNA phages, including CBASS and type I RM systems, both of which are widely distributed regardless of the pilin groups.

**Figure 5.**
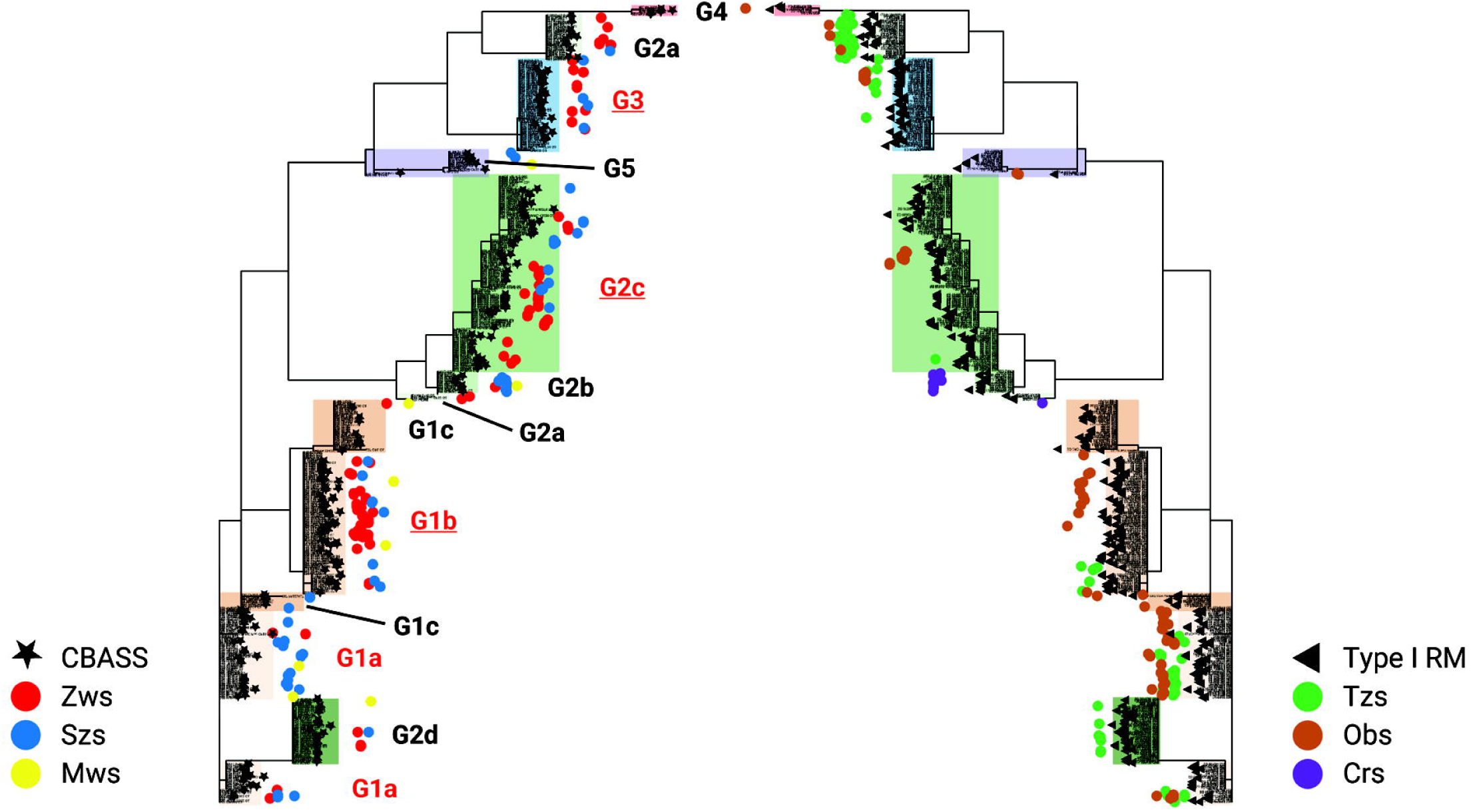
Co-occurrence of the pilin groups and the defense systems. The 653 complete genome sequences were retrieved from the *Pseudomonas* Genome Database (https://pseudomonas.com) and subjected to phylogenetic analysis based on the PilA sequences by using FastTree v.2.1.11, resulting in the phylogenetic tree generated by iTOL v.7. The genomes harboring the six RNA phage defense genes are designated as colored dots: *zws* (red), *szs* (blue), and *mws* (yellow) in the left; *tzs* (green), *obs* (brown), and *crs* (purple). CBASS (star) and type I RM (triangle) systems are also designated for comparison. The pilin groups susceptible to the RNA phages PP7 and LeivOr01 are highlighted in red color. Underlines indicate the pilin groups (G1b, G2c, and G3), where the Zws system is enriched.

Notably, Zws appears also enriched among strains carrying G3 pilins (11.9% vs. 16.8%), raising the possibility that G3 pilins may serve as adsorption scaffolds for RNA phages that have not yet been characterized. Together, these observations suggest that when infection exclusion is absent, acquisition of Zws-mediated intracellular defense may provide an additional layer of protection against RNA phage infection.

### ZwsA is an RNA endonuclease with selectivity against RNA phage genomes

Among the defense systems identified here, Zws was the most prevalent RNA phage-selective system and was therefore prioritized for mechanistic analysis. Its genetic organization and functional profile (Figure 4) suggest that the Zws system is a stand-alone defense effector (ZwsA) against RNA phages. As shown in Figure 6A and B, ZwsA contains VHS, DUF647, NERD, and S4 RNA-binding domains, with a putative catalytic pocket enriched in positive charge at the NERD domain. The NERD domain was originally described in nucleotide excision repair proteins such as UvrB and UvrC, where it has been linked to recognition of branched or damaged DNA structures [35]. Notably, NERD-domain proteins are frequently found within defense islands, although their roles in antiphage immunity remain poorly defined [28, 36].

**Figure 6.**
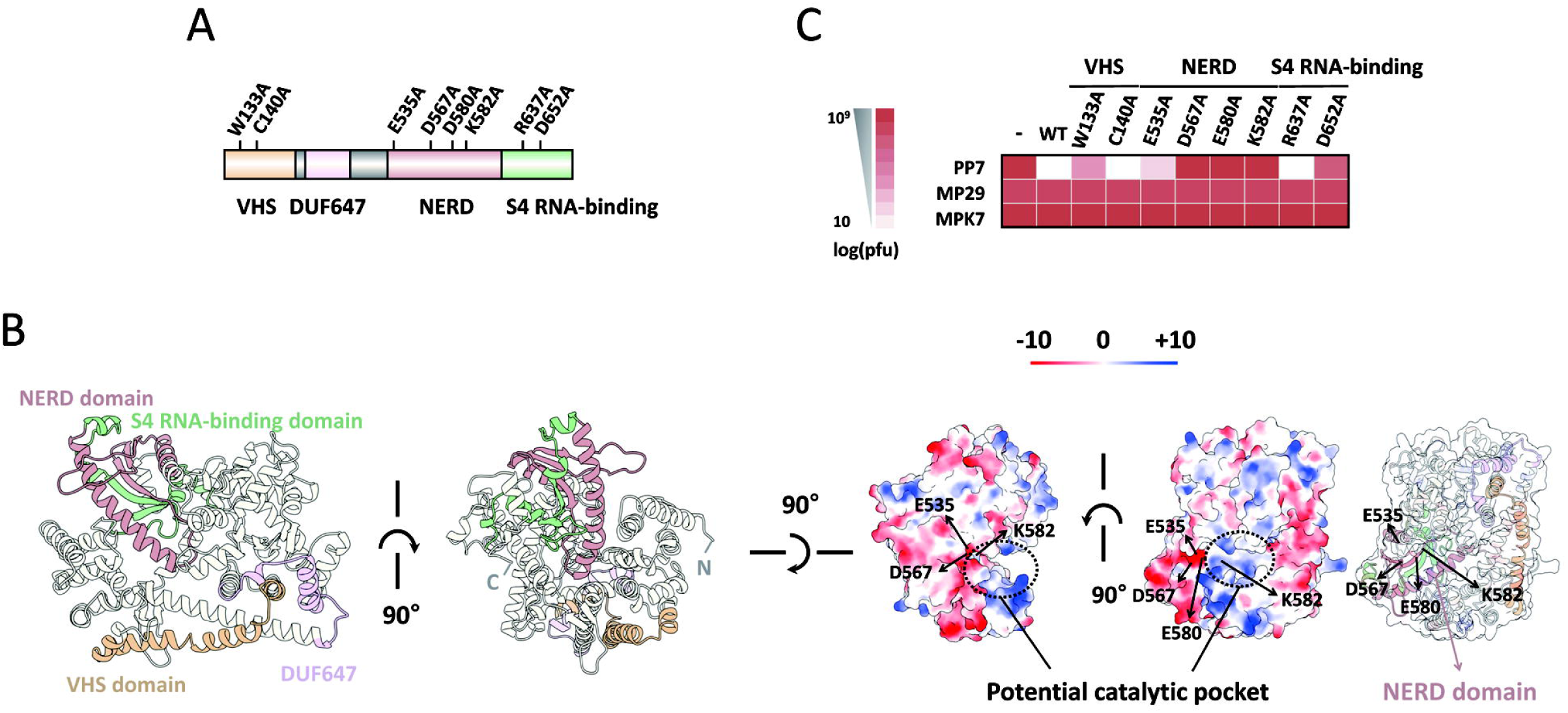
Structural modeling and domain function of ZwsA. **A**. Domain architecture of ZwsA and positions of the point mutants used for functional analysis in **C**. Predicted domains include VHS, DUF647, NERD, and an S4 RNA-binding domains. **B**. Structural modeling of ZwsA shown in multiple orientations, highlighting VHS, DUF647, NERD, and S4 RNA-binding domains, and a potential catalytic pocket centered with the positive charge (blue) within the NERD domain. Red designates the negatively-charged surface. The positions of the 4 amino acids (E535, D567, E580, and K582) are designated. **C**. Heat map to show the defense activity of the mutants in **A**. Efficiency of plaque formation (EOP) was assessed for PP7 and the DNA phages (MP29 and MPK7) as controls. Ten-fold serial dilutions of the phage lysates are spotted on a lawn of PAO1 cells with pUCP18 (-) or one of the pUCP18-derived plasmids for each mutant at VHS, NERD, and S4 RNA-binding domains. The color gradient represents the relative log(pfu) values. The values are the averages of three biological replicates.

To assess the roles of each domain of ZwsA in antiphage defense, we generated a panel of ZwsA point mutants (Figures 6A and C and S3). Four substitutions in the NERD domain, together with one substitution each in the VHS and S4 RNA-binding domains, impaired antiphage activity against PP7. This result indicates that multiple domains are required for full defense function, with the NERD domain likely serving a central role with the K582 residue within the catalytic pocket.

We next purified the ZwsA proteins to determine whether it functions as a stand-alone effector. ZwsA(K582A) was included as the inactive ZwsA control (Figure 7A). Using in vitro-transcribed PP7 genomic RNA, MS2 genomic RNA, and *lacZ* mRNA as substrates, we tested RNA cleavage activity in vitro. ZwsA cleaved both PP7 and MS2 RNA, whereas *lacZ* mRNA was unaffected (Figure 7B), indicating that ZwsA functions as an RNA endonuclease with substrate selectivity. Partial cleavage of PP7 and MS2 RNA was also observed with ZwsA(K582A), suggesting that this mutant retains residual activity. Nonetheless, neither wild-type nor mutant ZwsA cleaved *lacZ* mRNA, reinforcing the conclusion that ZwsA preferentially targets phage RNA rather than cellular mRNA.

**Figure 7.**
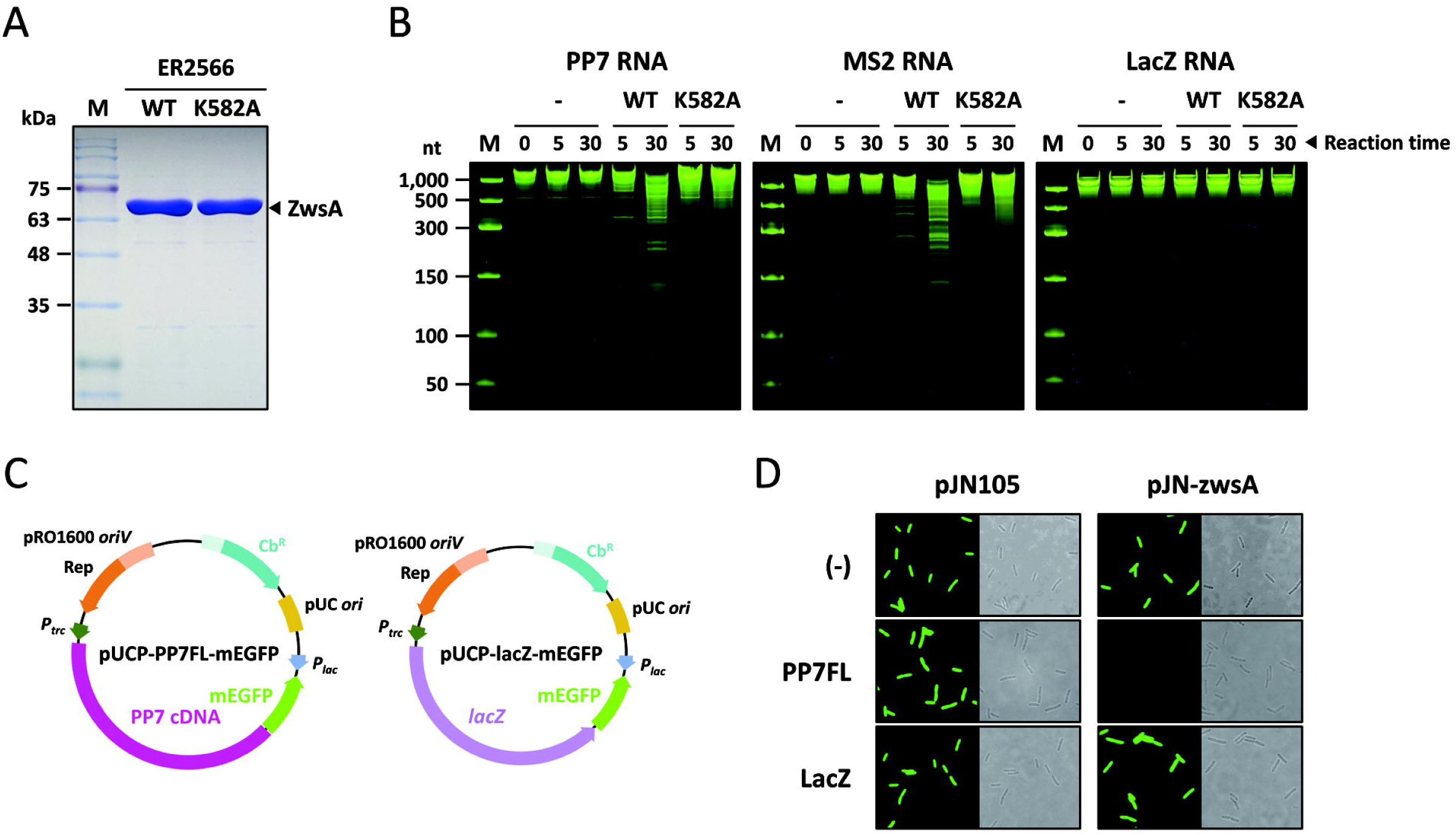
RNA endonuclease activity of ZwsA. **A**. ZwsA protein expression and purification. Heterologously expressed ZwsA (WT) and its mutant (K582A) proteins in *E. coli* ER2566 were analyzed by SDS-PAGE after affinity purification. Arrowhead indicates the ZwsA bands, whose deduced mass is 82.4 kDa. **B**. RNA cleavage activity of ZwsA in vitro. In vitro transcribed RNA samples of PP7, MS2 and *lacZ* were incubated for the designated time (0, 5, and 30 min) in the absence (-) or presence of the ZwsA proteins (WT and K582A) and then analyzed by Urea-PAGE. M represents RNA size marker for the designated sizes. **C**. Reporter plasmid maps for in vivo verification of ZwsA activity. pUCP18-based reporter systems based on mEGFP to generate the fused transcripts (PP7FL-mEGFP or lacZ-mEGFP) driven from the *P_trc_* promoter. **D**. Verification of RNA cleavage by ZwsA in vivo. The plasmids in **C** and their empty vector (-) are introduced into PAO1 cells with either the empty vector (pJN105) or the pJN105-borne *zwsA* (pJN-zwsA). The resulting 6 cells were grown for 3 h, placed on agarose pad, and observed by fluorescent microscopy. Representative images are shown with the corresponding bright field images.

To test this selectivity in vivo, we designed a reporter system to monitor ZwsA-dependent RNA cleavage (Figure 7C). pUCP18-based reporter plasmids were constructed to generate either the PP7 genomic RNA or the *lacZ* mRNA as a long transcript fused to the mEGFP mRNA. Because transcripts generated in vivo initially bear a 5′-triphosphate, they are relatively resistant to exonucleolytic degradation [37, 38]; endonucleolytic cleavage upstream of the mEGFP would instead generate a 5′-monophosphate and destabilize the mRNA. These reporter plasmids were co-introduced with a compatible ZwsA expression plasmid (pJN105), and mEGFP fluorescence was monitored by fluorescence microscopy (Figure 7D). Only the reporter containing PP7 cDNA lost fluorescence in the presence of ZwsA. Together, these findings support a model in which ZwsA acts as a stand-alone RNA endonuclease, likely through its NERD domain, to target RNA phage genomes while sparing host mRNAs. The precise RNA signature or structural feature recognized by ZwsA remains to be determined.

Transcriptome analysis further supported the selective nature of ZwsA activity. As shown in Figure S4, expression of ZwsA or SzsA alone did not substantially alter global transcriptomic profiles, whereas the presence of PP7 cDNA, leading to the increase in the PP7 RNA genome, induced clear transcriptional changes. Notably, transcriptomes associated with ZwsA and SzsA differed in the presence of PP7 cDNA, whereas inactive ZwsA did not measurably alter the transcriptomic response. These findings are consistent with ZwsA functioning as a bacterial defense effector that selectively targets phage RNA with minimal impact on host gene expression in the absence of RNA phage infection.

## DISCUSSION

This study expands the known repertoire of bacterial antiphage immunity by identifying six previously uncharacterized defense systems active against RNA phages. A key conceptual advance is that these systems were uncovered through a discovery strategy designed to interrogate intracellular immunity by bypassing phage adsorption and genome entry using cDNA-based RNA phage production. Because many bacterial strains often vary in phage-adsorbing scaffolds such as TFP and lipopolysaccharide for PA strains, conventional screens are frequently confounded by receptor-defect phenotypes that block infection at the cell surface and thereby mask intracellular defense mechanisms. Given that the infection exclusion involving receptor blockade is the prevalent defense systems in *E. coli* strains [39, 40], this approach provides a practical route for identifying intracellular defense genes independently of receptor variation by scoring cDNA-based RNA phage productivity. More importantly, this work not only reveals defense systems against RNA phages but also provides a broadly useful new ground for identifying new defense systems through the bioinformatic approaches based on guilt-by-association with these new systems.

The genomic analyses further place these systems within defense islands at specific genomic locations of the PA strains, emphasizing the coexistence of the known DNA phage defense systems such as Gabija, Zorya, Druantia, and restriction-modification modules. More importantly, the coexistence of multiple RNA phage defense systems has been functionally characterized in this study: in case of PMM44, three defense systems (Zws, Mws, and Crs), all of which are selective against RNA phages. The phenotypes of the isolated mutants suggested that they may be redundant, in that the deletion of one gene showed the maximal RNA phage productivity from the cDNA in these limited experimental conditions using one or two RNA phage strains. The presence of more than one RNA phage defense system in individual strains raises the possibility that combinatorial defense provides broader coverage, reduces escape, or balances protection with host fitness. One important implication for future research is how these multiple defense systems are coordinated to work, if they may partition their own roles across phage strains and/or infection stages. Therefore, it needs to be tested how multiple defense systems interact genetically and biochemically, including whether they are additive, hierarchical, or synergistic.

Our findings also point to an underappreciated principle in bacterial antiviral immunity: several of the systems identified here (Zws, Szs, Mws, and most likely Crs) acted selectively against RNA phages and did not protect against DNA phages tested. This pattern is consistent with the idea that distinct defense strategies have evolved against RNA and DNA phages, although weak or cryptic activities against RNA phages have been reported for some previously described systems such as Zorya and ApeA [41, 42]. The generality of this principle remains to be tested more broadly, because the present study was inevitably limited by the small number of available RNA phages. Even so, the selective activities observed here argue that RNA phages are not simply controlled collaterally by the DNA phage defense systems, but can instead be targeted by dedicated and mechanistically specialized defense modules. Although MS2 and PRR1 phages have been tested in our research, more RNA phages need to be characterized to elucidate the generality and specificity of the defense systems in this study. The co-occurrence of the susceptible pilins and the Zws system led us to hypothesize that the G3 pilins could be selected by the RNA phages yet-to-be discovered.

Biochemical and genetic analyses support ZwsA as a stand-alone defense effector that preferentially targets RNA substrates both in vitro and in vivo. This substrate selectivity is especially notable because it suggests a mechanism for discriminating phage genomic RNA from host cellular transcripts. One attractive possibility is that ZwsA recognizes higher-order structural signatures associated with RNA phage genomes, potentially including features linked to highly selective genome packaging [43, 44]. Although the precise recognition logic remains unresolved, defining the RNA features recognized by ZwsA and comparing them with those targeted by other RNA phage-selective systems such as Szs and Mws will be important for understanding how bacteria distinguish viral RNA from cellular RNA. More broadly, such knowledge could provide a conceptual bridge between bacterial antiviral immunity and structure-aware RNA biotechnology.

### Limitations of the study

This study has several limitations. First, we did not directly test defense activity in the native host backgrounds for all the identified defense systems, primarily because of twitching defects and/or pilin incompatibility with the limited RNA phage panel available. Second, although our data strongly support antiphage functions for these systems, additional work will be required to determine how each system is regulated in its native genomic context, particularly in the strains that encode multiple defense systems. Future studies should also address how these systems interact genetically and functionally during phage infection and whether they operate additively, hierarchically, or synergistically.

## ACKNOWLEDGMENTS

This work was supported by the National Research Foundation of Korea (NRF) Grants (RS-2022-NR070540, RS-2022-NR067344, and RS-2025-16069174)

## AUTHOR CONTRIBUTIONS

H.-W.B. and Y.-H.C. conceived and designed the research. H.-W.B., H.-J.K., and S.-Y.C. designed and performed the experiments and collected and analyzed the experimental data. H.-G.C., C.-H.W., M.-J.K., and H.-J.C. provided the reagents. H.-W.B. and Y.-H.C. wrote the manuscript. All authors reviewed the manuscript.

## METHODS

### Bacterial strains and growth conditions

All PA strains used in this study, including PAO1, PA14, PAK, the clinical and environmental isolates, and their derivatives, as well as the *Escherichia coli* strains DH5α, HB101, S17-1, SM10(λ*pir*), and ER2566, were routinely cultured at 37°C in Luria–Bertani (LB) broth or on LB plates containing 2% Bacto-agar (Difco). Selective cultivation of PA strains was performed on cetrimide agar (Difco) [17]. Where appropriate, antibiotics were added at the following final concentrations (μg/ml): gentamicin (50) or carbenicillin (200) for PA, and gentamicin (25) or ampicillin (50) for *E. coli*.

### Phage strains and lysate preparation

Phage lysates of MPK3, MPK7, MP29, and MP1413 were prepared by the plate lysate method using PAO1ΔRF as the host strain. By contrast, PP7 and LeviOr01 lysates were prepared from the supernatants of PAKΔRF cells carrying chromosomally integrated phage cDNA after growth in LB broth at 30°C for 24 h, as previously described [17, 23]. Phage particles were precipitated by adding 1 M NaCl and 10% polyethylene glycol 8,000 (Sigma-Aldrich), followed by incubation at 4°C overnight. The precipitated particles were collected by centrifugation and resuspended in 5 ml of phage buffer (50 mM Tris-HCl [pH 7.5], 10 mM MgSO_4_, 100 mM NaCl). Further concentration was achieved by ultracentrifugation at 180,000 × *g* for 6 h, after which the phage pellet was resuspended in 1 ml of phage buffer. The plaque forming units (pfu) of the phage lysates were determined before use.

### cDNA-based assembly of RNA phages in PA isolates

cDNA generated from phage-derived genomic RNA was cloned into a miniTn*7*-based vector and subsequently introduced into surrogate PA strains and a panel of 47 clinical and environmental isolates, as previously described [45]. Briefly, the cDNA was integrated chromosomally into PA strains by tetraparental conjugation involving two helper strains: a mobilizer strain carrying pRK2013 and a transposase donor strain carrying pTNS2 [45].

### Assessment of phage productivity

Phage productivity in PA strains carrying chromosomally integrated phage cDNA was assessed using plaque assay, spotting assay, and halo test using susceptible indicator strains. For plaque assays, appropriately diluted phage samples were mixed with 3 ml of 0.7% top agar containing ∼10^8^ late exponential phase indicator cells such as PAO1 lacking R and F pyocins (PAO1ΔRF), and overlaid onto LB agar plates. For halo tests, individual colonies of PA strains harboring PP7 cDNA was gently picked with a sterile toothpick and spotted onto lawns of PAO1ΔRF. For spotting assays, 3 μl of serially diluted phage samples prepared from the supernatants of PA strains carrying PP7 cDNA were spotted onto PAO1ΔRF lawns. Plates were incubated at 37°C for 16 h [17].

### Quantification of phage production

Phage production was quantified by plaque assay to determine pfu and by RT-qPCR following reverse transcription to enumerate RNA genome copy number, as previously described [45]. Phage genome copy number was measured using either phage lysates or phage-containing supernatants from cDNA-carrying PA strains. Phage RNA was extracted from phage samples using TRIzol (Invitrogen). To measure the intracellular RNA levels, cells were grown to mid-exponential growth phase and total RNA was extracted using the RNeasy Mini Kit (Qiagen). All RNA samples were treated with DNase I (Qiagen). cDNA was synthesized from 1 μg of RNA using the Toyobo ReverTra Ace qPCR RT kit and the PP7-RT-R primer. qPCR was then performed with the Toyobo Thunderbird SYBR qPCR mix and appropriate primers on a StepOnePlus real-time PCR system (Applied Biosystems). Relative phage genome copy numbers were normalized to the *rpoA* mRNA levels using standard curves. Phage copy number or virion counts were calculated as previously described [46].

### Transposon mutagenesis and screening

Random transposon mutagenesis was performed using the *mariner*-based transposon plasmid pBTK30 [47]. Biparental conjugation was carried out between SM10(λ*pir*) carrying pBTK30 and PA strains exhibiting reduced PP7 productivity on LB agar plates at 37°C for 6 h. Selection and counterselection of transposon insertion mutants were performed on cetrimide agar plates supplemented with gentamicin. In total, 140,869 insertion clones derived from 7 PA strains were screened initially by halo tests to identify mutants, in which halo size was restored to that observed in the reference strain. These primary candidates were then subjected to spotting assays for the semi-quantitative assessment of restored PP7 productivity. From this screen, 12 transposon mutants were selected, and the corresponding insertion sites were determined by arbitrary PCR followed by nucleotide sequencing, as described elsewhere [26].

### Heterologous expression of defense genes

The identified genes were amplified using gene-specific primers and cloned into the multicopy plasmid pUCP18. The resulting plasmids were introduced into the surrogate strains PAO1 and PMM3 by electroporation (Bio-Rad MicroPulser™) [46]. Defense activity in surrogate strains carrying individual pUCP18-borne defense genes was determined by spotting assay using a panel of RNA and DNA phages.

### Phylogenetic association of pilin groups and defense genes

PilA protein sequences were retrieved from 653 complete genomes deposited in the *Pseudomonas* Genome Database and subjected to multiple sequence alignments using MAFFT v.7.490. A phylogenetic tree was then generated from the aligned sequences using FastTree v.2.1.11 with default parameters. The resulting tree was exported in Newick format and visualized and annotated using iTOL v.7. Pilin groups were manually classified on the basis of phylogenetic clustering, and the associated defense systems among the 6 systems identified in this study were mapped on the phylogenetic tree.

### Generation of *zwsA* point mutants

Site-directed mutagenesis was used to generate alanine-substitution mutants in the VHS, NERD, and S4 RNA-binding domains of ZwsA. Mutations were introduced by four-primer splicing by overlap extension PCR, and the resulting amplicons were cloned into BamHI-digested pUCP18 by sequence- and ligation-independent cloning. The resulting plasmids were sequence-verified and introduced into PAO1 by electroporation. Defense activity was then evaluated by spotting assay using a panel of RNA and DNA phages.

### Structural modeling and analysis

Protein structure prediction was performed using AlphaFold3. The predicted ZwsA model was visualized and analyzed using UCSF ChimeraX (v.1.10rc). Domain architecture, including VHS, DUF647, NERD, and S4 RNA-binding domains, was annotated on the basis of sequence and structural features. Coulombic electrostatic surface potentials were calculated using UCSF ChimeraX and mapped onto the modeled protein surface. Structural representations were generated in multiple orientations, and a putative catalytic pocket was inferred from surface topology and charge distribution.

### Purification of ZwsA protein

ER2566 cells carrying pET15b-based constructs [48] encoding either wild type ZwsA or inactive ZwsA(K582A), each bearing N-terminal hexahistidine tag were, grown in LB broth at 37°C to mid exponential growth phase (OD_600_ = 0.2∼0.5). Cultures were then treated with 1 mM isopropyl-β-D-1-thiogalactopyranoside (IPTG) and incubated further overnight at 16°C. Cells were harvested, resuspended in 3 ml of binding buffer (50 mM Tris-HCl [pH 8.0], 300 mM NaCl, 5 mM imidazole), and lysed by sonication. Cell lysates were clarified by centrifugation at 13,523 × *g* for 10 min at 4°C. The supernatant was applied to 0.7 ml of TALON Superflow metal affinity resin (Takara) pre-equilibrated with binding buffer and incubated at 4°C for 1 h with rotation in a multimixer. The resin was washed 3 times with 4 ml of binding buffer, and bound proteins were eluted with elution buffer (50 mM Tris-HCl [pH 8.0], 300 mM NaCl, 200 mM imidazole). Protein purity was assessed by SDS-PAGE followed by Coomassie Brilliant Blue R staining. Samples were then concentrated using an Amicon centrifugal filter and used immediately for in vitro RNA cleavage assays.

### In vitro transcription of RNA substrates

RNA substrates were synthesized from the appropriate templates by in vitro transcription using the HiScribe T7 High Yield RNA Synthesis Kit (New England Biolabs). Briefly, DNA templates corresponding to PP7 cDNA, MS2 cDNA, and *lacZ* coding region, each placed downstream of a T7 promoter, were used for transcription. The resulting RNA transcripts were purified with the RNeasy Mini Kit (Qiagen). RNA integrity was assessed by electrophoresis on a 1% native agarose gel in TAE buffer (Tris-acetate-EDTA; 40 mM Tris, 20 mM acetic acid, 1 mM EDTA). Purified RNA samples were quantified before use.

### In vitro RNA cleavage assay

RNA cleavage assays were performed in 50 μl of reaction mixture, which contains 10 nM of each RNA substrate and 100 nM ZwsA, either WT or K582A, in reaction buffer (5 mM Tris-HCl [pH 7.5], 10 mM NaCl, 10 mM KCl, 2 mM MgCl₂, 0.2 mM DTT, and 0.2 units/μl RNase inhibitor). Reaction mixtures were incubated at 37°C for up to 30 min and the reaction was terminated by addition of quenching buffer (100 mM Tris-HCl [pH 7.5], 12.5 mM EDTA, 150 mM NaCl, 0.5 mg/ml proteinase K), followed by RNA cleanup using a Qiagen kit. Samples were then mixed with RNA loading dye (New England Biolabs), denatured at 95°C for 3 min, and resolved on 6% polyacrylamide gels containing 8 M urea. Gels were stained with SYBR Green II (Invitrogen).

### Fluorescence reporter assay

The fluorescence reporter system was constructed based on a pUCP18 plasmid harboring the constitutive *P_trc_* promoter and the mEGFP gene (Figure 7C). Either PP7 cDNA or the *lacZ* coding region was cloned downstream of the *P_trc_* promoter such that transcription produced either PP7 genomic RNA or lacZ mRNA as a long fusion transcript linked to mEGFP mRNA. Both reporter plasmids were co-introduced into PAO1 cells with either the compatible empty vector pJN105 or pJN105 carrying *zwsA* (pJN-zwsA). Cells were grown to an OD_600_ of ∼0.8, and ZwsA expression was induced with 0.2% arabinose. Cells were then mounted on 1% agar pads prepared in distilled water for imaging. Fluorescence images were acquired using a fluorescence microscope (Zeiss Axio Observer Z1). All images were processed using Zeiss ZEN software (v.2.6).

### Statistical analysis

Data are presented as mean ± standard deviation along with individual data points. Statistical analyses were performed using GraphPad Prism (version 8.4.3). Group comparisons were conducted using paired one-tailed *t*-tests, and 95% confidence intervals were determined as previously described [49]. Differences were considered statistically significant at *p* < 0.05.

