## Supplementary material for "cDNA-guided functional selection uncovers selective defense systems against RNA phages": SI Figures S1-4

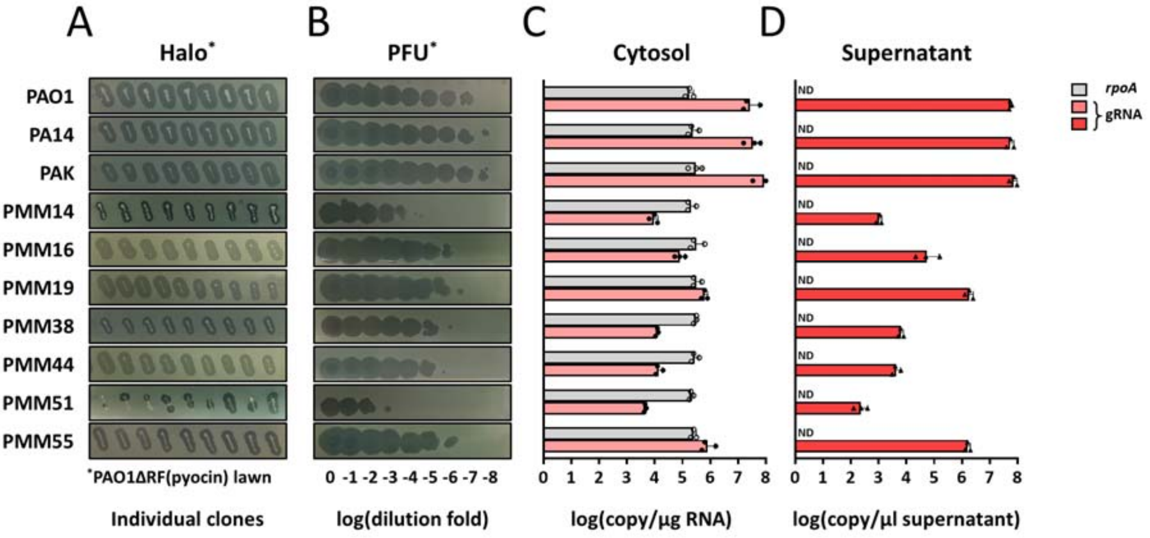


**Figure S1. Strains with reduced PP7 productivity**, related to Figures 1 and 2.

**A.** Halo test for PP7 progeny production from multiple individual clones of the 7 PMM strains identified as the PP7-reduced strains in this study. Control strains (PAO1, PA14, and PAK) with the PP7 cDNA were included for comparison. PAO1ΔRF was used as the susceptible lawn cells for PP7 infection.

**B.** Quantification of PP7 infectivity from a clone of the 7 PMM strains in **A**. Each culture supernatant was obtained from the 12 h broth cultures was subjected to 10 fold serial dilution for plaque formation unit (PFU) determination on PAO1ΔRF.

**C** and **D**. RT-qPCR analysis of PP7 genomic RNA (gRNA) in the cytosolic samples (pink bar) (**C**) and the culture supernatants (red bar) (**D**) from the clones of the PMM strains in **B**. The *rpoA* mRNA was used as the control (grey bar) as described in Methods. ND, not detected. Three independent biological replicates were used, with the individual data points and the error bars representing standard deviations.

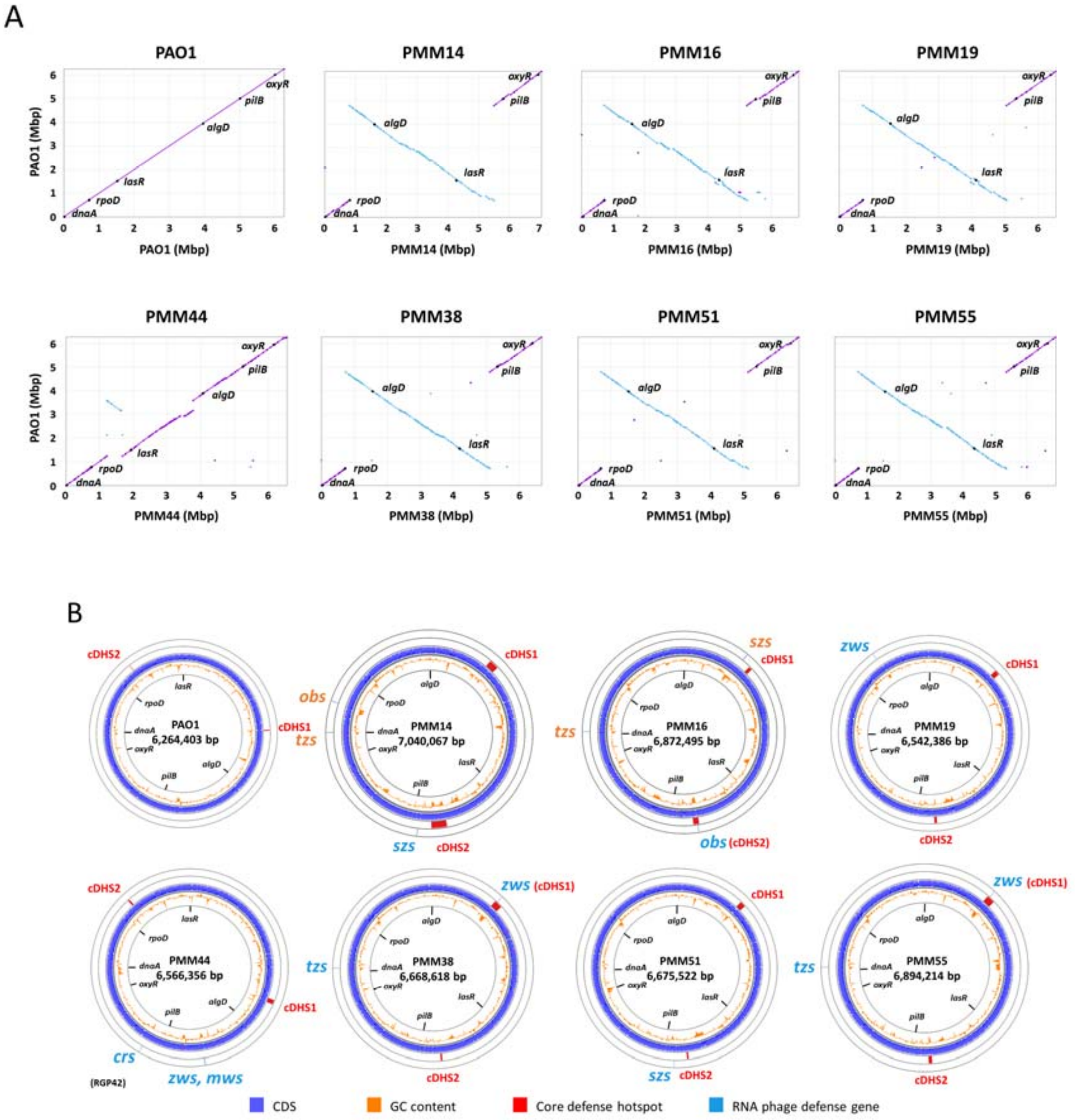


**Figure S2. Defense genes in the pangenome of PA strains with reduced PP7-productivity**, related to Figures 2 and 3.

**A**. Whole-genome synteny comparisons between PAO1 and the 7 PMM strains. The dot plots were generated based on the pair-wise genomic alignment by NUCmer (NUCleotide MUMmer) v3.1. The genomic coordinates are indicated by conserved chromosomal landmark genes in appropriate spacing (i.e. *dnaA*, *rpoD*, *lasR*, *algD*, *oxyR*, and *pilB*). Six strains have a large inversion as observed in PA14 (Lee et al. 2006).

**B**. Circular genome maps of the PMM strains showing the genomic locations of the 14 defense systems in Figure 3 against RNA phages. The maps were generated using Proksee (https://proksee.ca) (Grant et al. 2023), showing the GC content and core defense hotspots (cDHS1 and cDHS2) along each genome coordinate (Bae et al. 2025). The *oriC* regions of each genome are positioned at 9 o'clock and the clockwise direction follows the genome coordinate as in **A.**

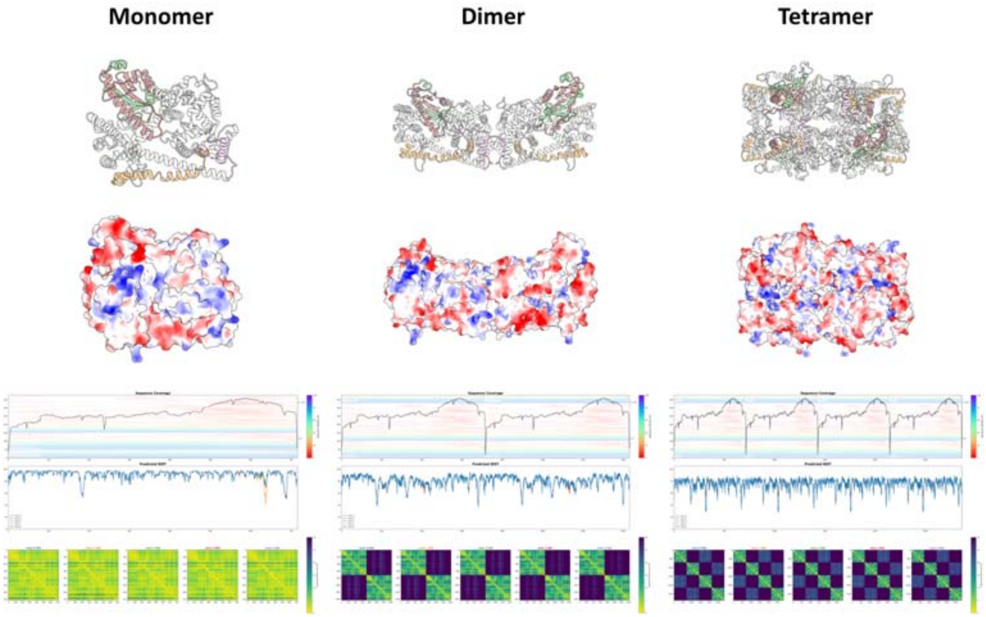


**Figure S3. Structural modeling of ZwsA**, related to Figure 6**.**

AlphaFold3-based structural models (top), electrostatic surface potentials (middle), and confidence metrics (bottom) for ZwsA monomer (left), dimer (center), and tetramer (right). Sequence coverage plots indicate the multiple sequence alignment (MSA) depth (black line, right log-scale y-axis), sequence identity (colored bars, rainbow scale), and unique sequence index (left y-axis) relative to the ZwsA query. The predicted local distance difference test (pLDDT) plots (left y-axis, 0–100) and the predicted alignment error (PAE) heatmaps (color scale 0–30 Å) reflect the confidences for top five ranked models, where lower PAE values (yellow) signify higher accuracy in intra- (diagonal) and inter-subunit (off-diagonal) alignment across the 716-residue subunits.

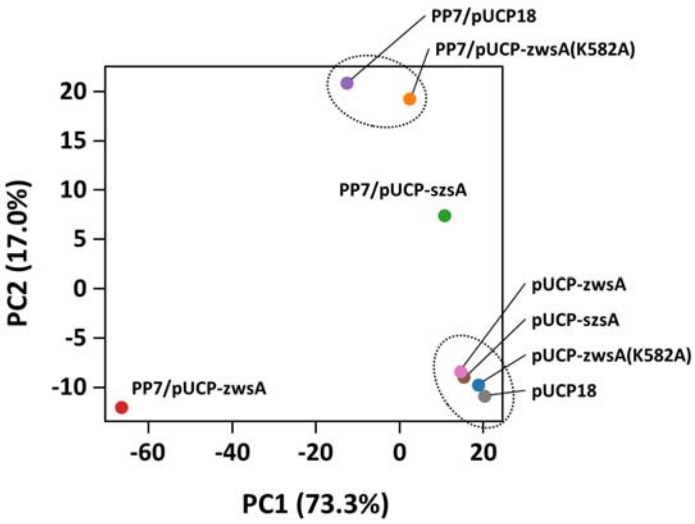


**Figure S4. Transcriptomic changes by ZwsA and SzsA**, related to Figure 7.

Multidimensional scaling plot of transcriptome profiles from PAO1 cells with or without the PP7 cDNA. The cells carry one of the 4 plasmids: empty vector (pUCP18) or one of the pUCP18 derivatives containing wild type ZwsA (pUCP-zwsA), the ZwsA catalytic mutant (pUCP-zwsA(K582A)), or wild type SzsA (pUCP-szsA). The 4 cell types without the PP7 cDNA were clustered (right bottom), irrespective of the carried plasmids, whereas the PP7-containing cells were scattered, except for the clustered 2 cell types that carry either pUCP18 or pUCP-zwsA(582A).
